# Deciphering the metabolic details of L-lysine toxicity in cyanobacteria

**DOI:** 10.1101/2025.03.10.642250

**Authors:** Andreas Enkerlin, Ute A. Hoffmann, Johanna Rapp, Rui Miao, Hannes Link, Elton P. Hudson, Khaled A. Selim

## Abstract

L-lysine (Lys) has been explored as a potential cyanobactericide due to its inhibitory effects on cyanobacterial growth at micromolar concentrations, comparable to many antibiotics. Here, we investigated the early metabolic and physiological responses of the model cyanobacterium *Synechocystis* sp. PCC 6803 to Lys exposure. Physiological analyses revealed cell enlargement, oxidative stress, and photosynthesis inhibition, leading to growth arrest. Metabolomic profiling indicated disruptions in peptidoglycan biosynthesis, evidenced by the accumulation of L-/D-alanine, meso-diaminopimelate, and D-Ala-D-Ala, suggesting interference with cell wall integrity. Furthermore, levels of energy metabolites and other amino acids including tyrosine, tryptophan, valine, and iso-/leucine were significantly altered, implying broader metabolic impacts of Lys toxicity. To explore potential resistance mechanisms, we used a CRISPRi-based genetic screen to identify key genes involved in relieving Lys toxicity. The Bgt permease system, responsible for basic amino acid uptake, was essential for acquiring Lys-resistance, as a *bgtA* mutant exhibited a normal growth on elevated Lys concentrations, thereby validating our CRISPRi-screen. Additionally, UirR, a DNA-binding response regulator, and genes linked to c-di-AMP signaling, seemed implicated in Lys metabolism. Deletion of c-di-AMP synthase gene increased Lys sensitivity, supporting a role for c-di-AMP in cell wall homeostasis and osmotic stress regulation. Altogether, our findings explored the early metabolic responses and physiological consequences of Lys exposure in *Synechocystis*, demonstrating its effects on peptidoglycan biosynthesis, amino acid metabolism, and nucleotide biosynthesis. The identification of key genetic factors contributing to Lys resistance provides new insights into cyanobacterial physiology and the potential application of Lys in bloom control strategies.

## 1. Introduction

Because of its micromolar inhibitory concentration and its importance in aquatic ecosystems as an essential amino acid for heterotrophs, L-lysine (Lys) has been investigated as a potential cyanobactericide for controlling cyanobacterial blooms since the early 21^st^ century (Hehmann *et al*., 2002; Takamura *et al*., 2004; Dahedl *et al*., 2023). Such blooms pose a threat to aquatic ecosystems and human health (Plaas & Paerl, 2021). Lys toxicity has been reported for some cyanobacteria, in particular for *Microcystis* sp. (Zimba *et al*., 2019; Takamura *et al*. 2004). For *Microcystis aeruginosa*, known for bloom formations and production of the microcystin toxin, Lys was reported to inhibit growth at as low as 34 µM and reduce chlorophyll A content (Tian *et al*., 2018). In an untargeted metabolomic analysis, Yan *et al*. (2023) found that after three days of Lys-treatment, *M. aeruginosa* amino acid metabolism was altered, and especially metabolites related to arginine and proline metabolism increased. After 30 days of treatment, in a targeted approach, they showed that out of 25 amino acids analyzed, 22 were significantly decreased, whereas L-arginine, L-ornithine, and L-citrulline showed a significant increment. Thus, they proposed that Lys inhibits arginine dihydrolase, leading to an accumulation of metabolites from the ornithine-ammonia (OA) cycle (Zhang *et al*., 2018). Recently, another study attributed cell death of *M. aeruginosa* by Lys substitution of meso-diaminopimelic acid (mDAP) at the peptidoglycan peptide chain, inhibiting cross-linkage and therefore interrupting peptidoglycan synthesis (Kim *et al*., 2023). Notably, the degree of cross-linkage (56%) in cyanobacteria, particularly *Synechocystis* sp. PCC 6803 (hereafter *Synechocystis*), falls closer in the range of Gram-positive bacteria, but its peptide bridges have a Gram-negative composition. The peptides are composed of L-alanine, D-glutamate, mDAP, and D-alanine, which are typical of Gram-negative bacteria, whereas in Gram-positive bacteria the third incorporated amino acid is Lys instead of mDAP (Jürgen *et al*., 1983).

Similar to *M. aeruginosa*, the growth of *Synechocystis* is inhibited at 33 µM Lys (Flores & Muro-Pastor, 1990). Though degradation and biosynthesis of Lys has not been studied in this model organism, its genome includes the mDAP pathway enzymes required for Lys biosynthesis (Hudson *et al*., 2006; Hudson *et al*., 2008). This pathway uses aspartate-4-semialdehyde as a substrate, which is derived from aspartate. In the final step, mDAP is converted to Lys by the diaminopimelate decarboxylase *lysA* (encoded by *sll0504*). Interestingly, in *Synechocystis* and many cyanobacteria, *lysA* forms a conserved operon with the c-di-AMP synthesis gene *dacA* (encoded by *sll0505*) and the undecaprenyl pyrophosphate synthase *uppS* (encoded by *sll0506*) that produces a lipid carrier for peptidoglycan synthesis (Selim *et al*. 2021). As all three genes are co-transcribed, this potentially implicates a role for c-di-AMP in cell wall homeostasis (Agostoni *et al*., 2018; Selim *et al*. 2021; Mantovani *et al*. 2023). On the catabolic side, *Synechocystis* genome contains a putative lysine decarboxylase *cad* (encoded by *sll1683*) (Kaneko *et al*., 1996) and an L-amino acid dehydrogenase capable of oxidizing Lys (Schriek *et al*., 2009).

As an autotroph, *Synechocystis* can synthesize all its essential amino acids from inorganic nitrogen and carbon sources using energy from photosynthesis. Nevertheless, it possesses multiple characterized and uncharacterized amino acid permeases (Quintero *et al*., 2001). Among a diverse range of tested cyanobacteria, *Synechocystis* showed the highest uptake of the basic amino acids L-arginine, Lys, and L-histidine (Montesinos *et al*., 1997). These amino acids, along with L-glutamine, are transported by the “basic amino acid and glutamine transporter” Bgt system encoded by *bgtA* (*slr1735*) and *bgtB* (*sll1270*), which are homologs of GlnQ and GlnH of the *Escherichia coli* glutamine ABC transporter, respectively. The BgtAB transporter was shown to have a high affinity for L-arginine, Lys, and L-histidine, and contrarily a low affinity and high capacity for L-glutamine (Labarre *et al*., 1987; Flores & Muro-Pastor, 1990; Quintero *et al*., 2001). Moreover, independent disruption of both subunits similarly reduced the transport of the four amino acids (Quintero *et al*., 2001).

The mechanisms behind Lys toxicity in *Synechocystis* have not been thoroughly studied before, yet it shows comparable sensitivity to *M. aeruginosa* and is a convenient model to study amino acid metabolism in cyanobacteria due to its faster growth rates and relatively simpler physiology. Here, we covered an early metabolic response of Lys toxicity on *Synechocystis* cells. We found that Lys mainly affected amino acid metabolism, primarily those related to peptidoglycan synthesis such as D-/L-alanine and mDAP. Our results support the notion of peptidoglycan crosslinking inhibition by Lys as observed in *M. aeruginosa*. Moreover, to understand the feasibility of utilizing Lys as a cyanobactericide against harmful blooms, we explored how the cells could acquire resistance against Lys toxicity using the advantage of pooled CRISPRi library screen (Yao *et al*., 2020; Miao & Jahn *et al*., 2023).

## 2. Results

### 2.1 Physiological response to lysine

To investigate the physiological response of *Synechocystis* under toxic Lys concentrations, we first determined the Lys inhibitory concentration by titrating a range of concentrations in well-plates. At 50 µM added Lys, no growth was observed (**Suppl. Fig. S1**), which is in accordance with 33 µM reported by (Flores & Muro-Pastor, 1990). At 25 µM, the growth was inhibited for 48 h and then resumed slowly after 120 h. Remarkably, cells were also able to recover from N-starvation induced chlorosis using this concentration and up to 75 µM, indicating that Lys is taken up and metabolized as a nitrogen source (**Suppl. Fig. S1**). Therefore, we chose 25 µM for the physiological assessment of Lys toxicity, as this concentration showed strong growth impairment but did not kill the cells.

Next, wild type (WT) *Synechocystis* cells were grown in BG11^N^ treated with and without 25 µM Lys. As expected, Lys-treated cells initially showed strong growth inhibition and resumed growing after 72-120 h (**Fig. 1A**). As an indication of the overall cell fitness and photosynthetic activity, we measured the apparent photosystem II (PSII) quantum yield using pulse-amplitude modulation (PAM) fluorescence. In accordance with the growth, the PSII activity of Lys-treated cells vanished almost to zero after 48 h and regained slowly after 72 h, while the PSII activity of untreated cells stayed almost constant (**Fig. 1B**). Similarly, the whole-cell spectrum analysis revealed a strong decrease in the photosynthetic pigmentation (phycobilisomes and chlorophyll a) under Lys-treatment compared to untreated cells (**Fig. 1C**).

**Figure 1.**
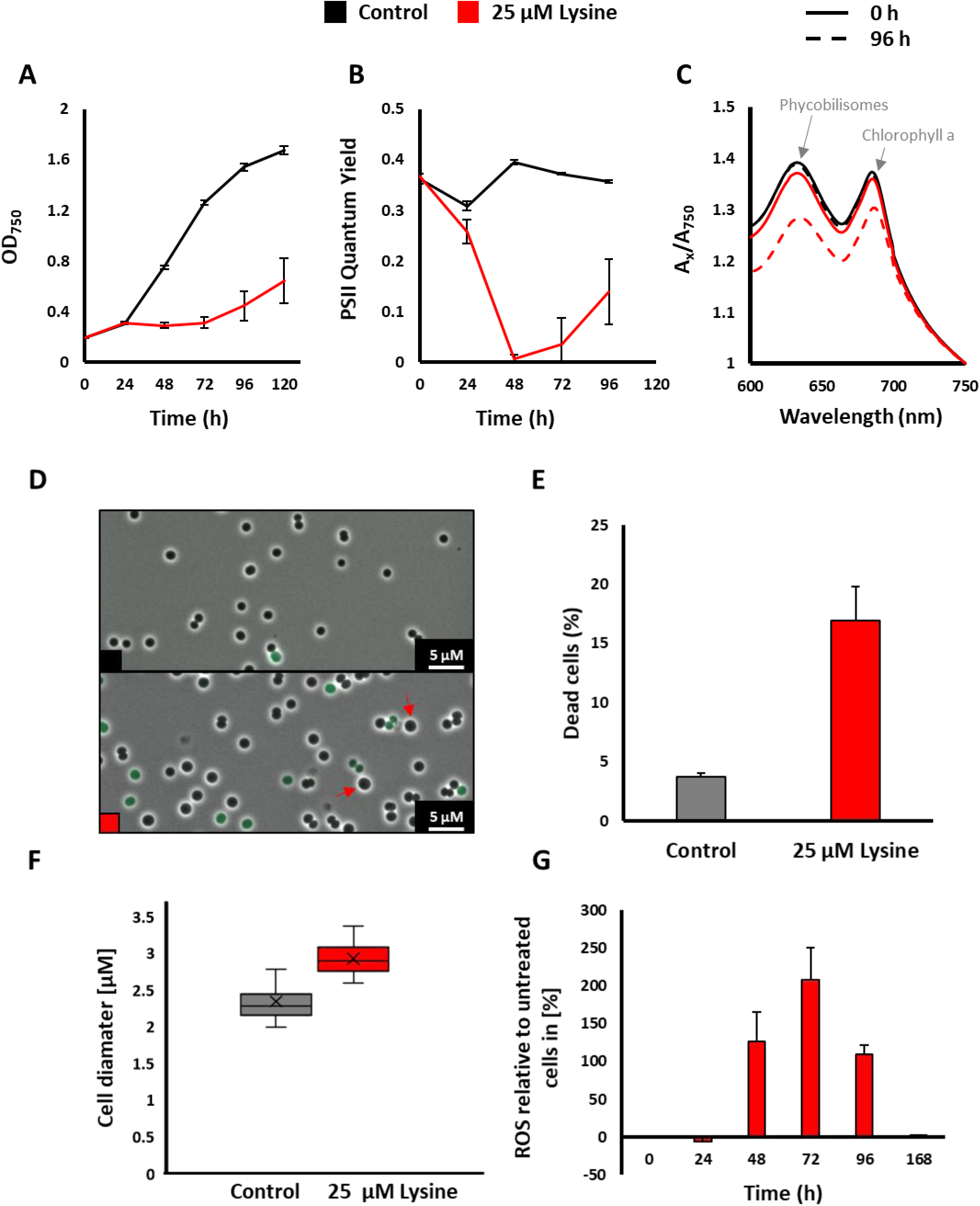
Physiological characterization of *Synechocystis* under Lys treatment with and without 25 µM Lys. **A:** Growth represented by the mean of OD_750_ ± SD from biological triplicates. **B**: photosynthetic efficiency measured by PAM fluorometer and showing the PSII quantum yield ± SD from biological triplicates. **C:** The whole-cell spectrum (WCS) of one replicate is shown at time points (0 and 96 h) of Lys-treatment. Absorbance values are normalized to OD 1.0 at 750 nm. Absorption peaks characteristic of phycobilisomes and chlorophyll a are indicated. **D:** Representative of DiBAC_4_(3) staining of cells after 48 h of Lys treatment. Green stained cells are dead cells. Some enlarged cells are indicated with arrows. **E:** Quantification of average dead cells (in percentage) stained with DiBAC_4_(3) from 500-1000 cells. **F:** Quantification of average cell diameter from 500-1000 cells. Bar and whisker denote 5^th^, 25^th^, median, 75^th^, and 95^th^ percentiles. **G:** Levels of reactive oxygen species (ROS) in Lys-treated cells relative to untreated cells (at time point zero). The presence of ROS was quantified using the H2DCFDA fluorescent marker (Haffner *et al*. 2023).

To find out whether Lys has a bactericidal effect at 25 µM Lys, fixed cells from 48 h of Lys-treatment were stained with DiBAC_4_(3), a dye that penetrates depolarized (*i*.*e*., dead) cells (**Fig. 1D**). Approximately 17-20% of treated cells were dead compared to ≈ 4% of untreated cells (**Fig. 1E**). Moreover, treated cells appeared enlarged, showing a 25 % increase in the average cell diameter (**Fig. 1F and Suppl. Fig. S2**), implying that Lys could potentially influence on the cell size and cell wall, or osmoregulation and maintenance of the intracellular turgor pressure.

### 2.2 Metabolic response to lysine

To assess the metabolic response accomplished with Lys stress, we performed targeted metabolomics via LC-MS/MS for carbon, nitrogen, nucleotide, and cell wall metabolism in the early stages of Lys-treatment (after 24 h, 48 h, 72 h, and 120 h of 25 µM Lys addition), as the physiological effect was already noticeable in the first 24 h (**Fig. 1B**). The measured average foldchanges are provided (**Table 1**).

**Table 1:** Heatmap of Log_2_(foldchanges) of metabolites measured by LC-MS/MS under 25 µM Lys-treatment. The foldchanges were calculated from relative concentrations to untreated samples (0 h), and the log_2_ of the normalized values are shown in the heatmap with red being upregulated and blue downregulated.

| Metabolites | Log<sub>2</sub>(Foldchange) |  |  |  |  |  | Foldchange |
| --- | --- | --- | --- | --- | --- | --- | --- |
|  | 0 h | 24 h | 48 h | 72 h | 120 h | AVG (24-120 h) | AVG (24-120 h) |
| Alanine |  |  |  |  |  | 4.63 | 24.77 |
| Lysine |  |  |  |  |  | 4.59 | 24.08 |
| D-Pantothenic acid (CoA) |  |  |  |  |  | 3.70 | 13.02 |
| Serine |  |  |  |  |  | 3.10 | 8.57 |
| S-Adenosylmethionine |  |  |  |  |  | 2.92 | 7.56 |
| Diaminopimelic acid (mDAP) |  |  |  |  |  | 2.91 | 7.54 |
| D-Alanyl-Alanine |  |  |  |  |  | 2.90 | 7.48 |
| Tryptophan |  |  |  |  |  | 2.88 | 7.34 |
| Oxidized Glutathione |  |  |  |  |  | 2.81 | 7.00 |
| Valine |  |  |  |  |  | 2.78 | 6.87 |
| Guanosine |  |  |  |  |  | 2.71 | 6.55 |
| Tyrosine |  |  |  |  |  | 2.68 | 6.40 |
| Adenosine |  |  |  |  |  | 2.42 | 5.36 |
| Iso-/leucine |  |  |  |  |  | 2.08 | 4.22 |
| Cytidine |  |  |  |  |  | 1.86 | 3.63 |
| Citrulline |  |  |  |  |  | 1.77 | 3.42 |
| Guanine |  |  |  |  |  | 1.70 | 3.24 |
| N2-Succinyl-L-arginine |  |  |  |  |  | 1.68 | 3.20 |
| N-Acetyl-L-Ornithine |  |  |  |  |  | 1.64 | 3.12 |
| Glycine |  |  |  |  |  | 1.64 | 3.12 |
| N-Acetyl-L-Glutamic acid |  |  |  |  |  | 1.49 | 2.80 |
| Hexose-P |  |  |  |  |  | 1.46 | 2.75 |
| CMP |  |  |  |  |  | 1.19 | 2.28 |
| GMP |  |  |  |  |  | 1.16 | 2.23 |
| Glutamic acid |  |  |  |  |  | 1.14 | 2.21 |
| Adenine |  |  |  |  |  | 1.05 | 2.06 |
| Histidine |  |  |  |  |  | 1.02 | 2.03 |
| Arginine |  |  |  |  |  | 0.93 | 1.90 |
| AMP |  |  |  |  |  | 0.89 | 1.85 |
| Methionine |  |  |  |  |  | 0.84 | 1.78 |
| Adenylosuccinic acid |  |  |  |  |  | 0.77 | 1.71 |
| UDP-N-acetyl-D-Glucosamine |  |  |  |  |  | 0.75 | 1.69 |
| Proline |  |  |  |  |  | 0.65 | 1.57 |
| Glutamine |  |  |  |  |  | 0.61 | 1.52 |
| Dihydroxyacetone-P |  |  |  |  |  | 0.57 | 1.49 |
| ADP |  |  |  |  |  | 0.57 | 1.48 |
| Succinate |  |  |  |  |  | 0.52 | 1.43 |
| Glycerol 3-Phosphate |  |  |  |  |  | 0.50 | 1.42 |
| NADP |  |  |  |  |  | 0.46 | 1.38 |
| IMP |  |  |  |  |  | 0.40 | 1.32 |
| Pyridoxamine 5-Phosphate |  |  |  |  |  | 0.35 | 1.27 |
| S-Adenosyl-L-homocysteine |  |  |  |  |  | 0.29 | 1.22 |
| GABA |  |  |  |  |  | 0.27 | 1.21 |
| Ornithine |  |  |  |  |  | 0.25 | 1.19 |
| Protoheme |  |  |  |  |  | 0.21 | 1.16 |
| ATP |  |  |  |  |  | 0.18 | 1.13 |
| Aspartic acid |  |  |  |  |  | 0.16 | 1.12 |
| NAD |  |  |  |  |  | 0.14 | 1.10 |
| N-Carbamoyl-L-aspartate |  |  |  |  |  | 0.13 | 1.10 |
| Acetyl-CoA (AcCoA) |  |  |  |  |  | 0.12 | 1.08 |
| Flavin adenine dinucleotide |  |  |  |  |  | 0.04 | 1.03 |
| Asparagine |  |  |  |  |  | -0.03 | 0.98 |
| 2-Deoxyadenosine |  |  |  |  |  | -0.07 | 0.95 |
| UMP |  |  |  |  |  | -0.10 | 0.93 |
| Urea (Carbamide) |  |  |  |  |  | -0.14 | 0.91 |
| Phenylalanine |  |  |  |  |  | -0.18 | 0.88 |
| 7,8-Diaminononanoate |  |  |  |  |  | -0.20 | 0.87 |
| Citrate |  |  |  |  |  | -0.21 | 0.86 |
| UDP-Glucose |  |  |  |  |  | -0.24 | 0.85 |
| Riboflavin |  |  |  |  |  | -0.65 | 0.64 |
| Phosphoenolpyruvate |  |  |  |  |  | -0.88 | 0.54 |
| Propionyl-CoA |  |  |  |  |  | -0.89 | 0.54 |
| Dihydropteroate |  |  |  |  |  | -2.83 | 0.14 |

#### 2.2.1 Cell wall- and amino acid metabolism

Due to its involvement in the mDAP pathway as well as its possible integration into the cell wall (Kim *et al*., 2023), we hypothesized that Lys would disturb the homeostasis of metabolites related to cell wall synthesis. **Figure 2** depicts a schematic of central metabolism and the relative Log_2_(foldchanges). Among all analyzed metabolites, Lys showed the second highest average foldchange (24.08; **Table 1**), indicating that it was taken up into the cells. Although the foldchange gradually decreased after 48 h, the lowest foldchange (observed at 120 h) was 13.06 (**Table 1**), suggesting that Lys was metabolized slowly. *Synechocystis* contains the complete mDAP pathway for Lys biosynthesis, where mDAP is converted into Lys by the mDAP decarboxylase (*lysA*) in the final step (Mills *et al*., 2020). In accordance, mDAP was also among the highest average foldchanges with 7.54 (**Table 1**). In *Synechocystis*, mDAP is the third incorporated amino acid in the peptidoglycan peptide bridge. These peptides are composed of L-alanine, D-glutamate, mDAP, and D-alanine (Jürgen *et al*., 1983). Interestingly, alanine had the highest average foldchange of 24.77 out of all the metabolites analyzed (**Table 1**; **Fig. 2**), implying that the alanine was not incorporated to the growing peptide chain. During transpeptidation, the dipeptide D-alanyl-D-alanine forms in the terminal end of the peptide bridge before cleavage of one of the D-alanines. D-alanyl-D-alanine had the seventh highest average foldchange of 7.48 (**Table 1**), further supporting the dysregulation of transpeptidation of growing peptidoglycan after Lys-treatment. The accumulation of mDAP, alanine and D-alanyl-D-alanine are in accordance with the recent finding of mDAP being outcompeted by the high levels of Lys (**Table 1 and Fig. 2**), leading to false incorporation of Lys instead of mDAP into peptidoglycan by an unspecific UDP-N-acetylmuramoyl-L-alanyl-D-glutamate-2,6-diaminopimelate (MurE) ligase (Kim *et al*. 2023). *Synechocystis* MurE shares high degree of identity to *M. aeruginosa* MurE (**Suppl. Fig. S3**) and therefore likely to falsely incorporate Lys into peptidoglycan, which could explain the overaccumulation of these metabolites, and thus the subsequent growth arrest due to the inhibition of transpeptidation. This result is also in agreement with enlargement of cells after Lys-treatment (**Fig. 1F**), likely due to cell wall weakening and impaired crosslinking via Lys misincorporation into the peptide bridges (Kim *et al*., 2023).

**Figure 2.**
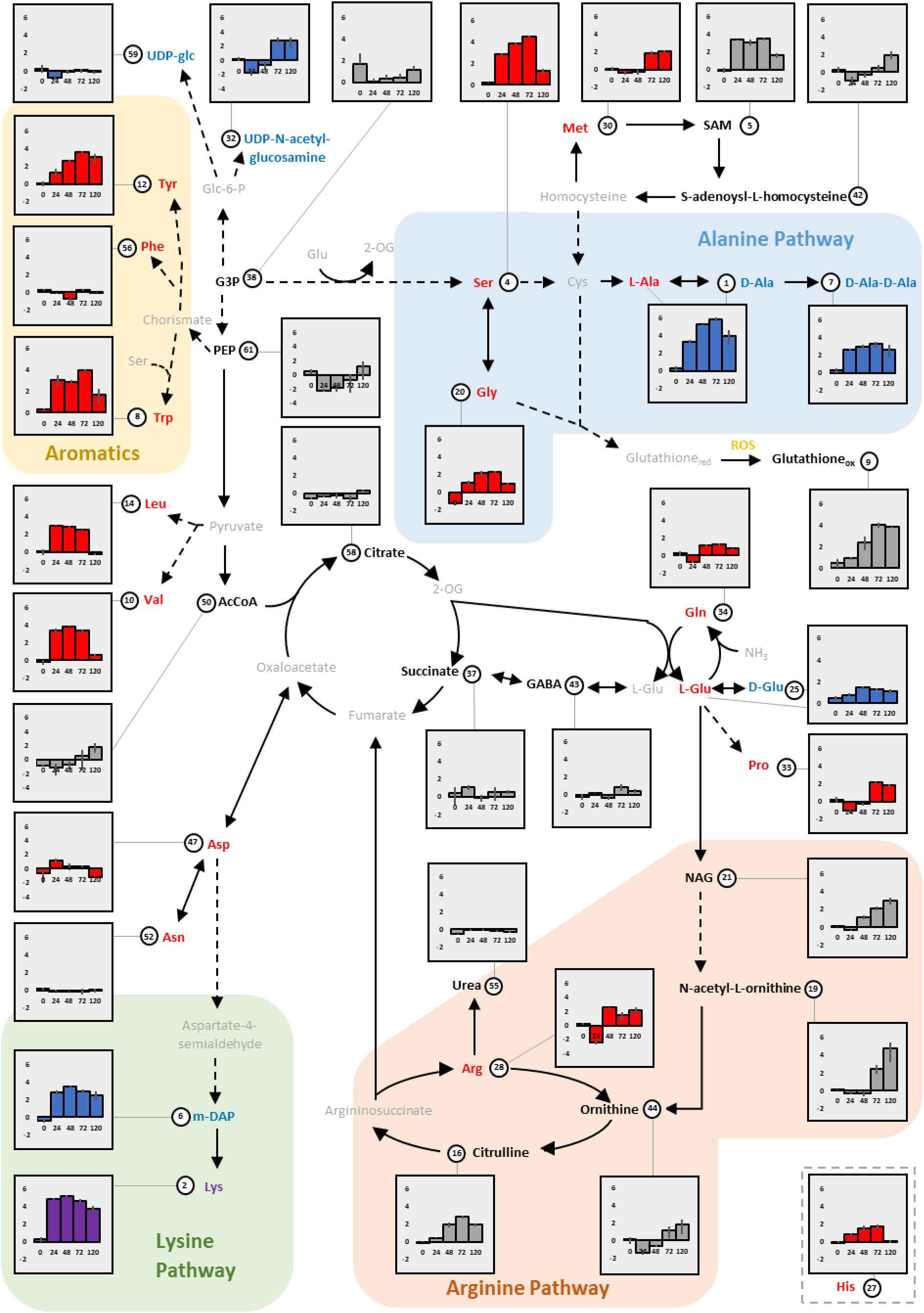
LC-MS/MS analysis of metabolome of *Synechocystis* WT grown in BG11^N^ with or without 25 µM L-lysine (Lys). Metabolites were quantified before addition of Lys (0 h) and at four different time points after Lys-treatment (24, 48, 72 and 120 h). The graphs represent the Log_2_(foldchanges) of Lys-treated cells relative to untreated cells. The error bars represent the standard deviation of Log_2_(foldchange) between treated and untreated cells. The pathways for alanine, arginine, and Lys are highlighted, as well as the aromatic amino acids. Lys is written in **purple**, cell wall-related metabolites in **blue**, proteinogenic amino acids in **red**, and other measured metabolites in **black**. **ROS** = reactive oxygen species. X-axis: Time (h). Y-axis: relative Log_2_(foldchange) between Lys-treated and untreated cells. Error bars represent the standard deviations of treated cells compared to the means of untreated cells.

Alanine biosynthesis includes the two-step conversion of L-serine into L-cysteine and a subsequent desulfonation into L-alanine. Finally, the interconversion of L-alanine enantiomer to D-alanine, a constituent of peptidoglycan (Mills *et al*., 2020), is catalyzed by the alanine racemase *alaR* (encoded by *slr0823*) (Ashida *et al*., 2022). Although cysteine was not quantified, *Synechocystis* possesses putative enzymes that catalyze the conversion of homocysteine to L-methionine and then to S-adenosyl-L-methionine (SAM), which reported the fifth highest average foldchange of 7.56 (**Table 1**) (Mills *et al*., 2020). Moreover, the accumulation of alanine has a negative feedback effect on alanine synthesis, leading to accumulation of serine. Thus, serine showed the fourth highest average foldchange of 8.57 (**Fig. 2**). Alternatively, L-serine can be synthesized via the glyoxylate pathway from L-glycine, which was among the upregulated metabolites (**Table 1**). Serine is also required for the synthesis of tryptophan, which had the eighth highest average foldchange of 7.34 (**Table 1**).

The other aromatic amino acids tyrosine and phenylalanine had average foldchanges of 6.40 and 0.88, respectively (**Table 1**; **Fig. 2**). Besides alanine and mDAP, other metabolites related to cell wall metabolism such as UDP-glc (average foldchange: 0.85), UDP-N-acetyl-glucosamine (average foldchange: 1.69), and (D-)glutamate (average foldchange: 2.21) were slightly upregulated. Importantly, the peptidoglycan precursor UDP-N-acetyl-glucosamine decreased in the first 24 h relative to the untreated cells and later increased (**Table 1**), further supporting the notion of impaired peptidoglycan biosynthesis. Intriguingly, amino acids derived from L-glutamate such as L-glutamine, L-proline, L-arginine, L-ornithine, citrulline, N-acetyl-L-ornithine, and N-acetyl-L-glutamic acid (NAG) did not show high foldchanges compared to those from the alanine and lysine pathways (**Table 1**), but instead decreased the first 24 h and increased slightly after 24-48 h (**Table 1 and Fig. 2**). Other amino acids that reported high average foldchanges were valine (6.87) and iso-/leucine (4.22) (**Table 1**). Notably, our results revealed a similar trend like what has been reported for *M. aeruginosa* with an upregulation of arginine and proline metabolism after 72 h of Lys-treatment. The long-term Lys-treatment of *M. aeruginosa* led additionally to accumulation of the ornithine-ammonia (OA) cycle metabolites (arginine, ornithine, and citrulline) (Yan *et al*., 2023).

#### 2.2.2 Carbon-, nucleotide-, and oxidative stress metabolisms

In contrast to amino acids and nitrogen metabolism, carbon metabolism of Lys-treated cells did not differ much from untreated cells, as exemplified by the average foldchanges from metabolites of the tricarboxylic acid (TCA) cycle such as citrate (0.86) and succinate (1.43) (**Table 1**). However, the fatty-acid metabolites (propionyl-CoA and acetyl-CoA) and the phosphoenolpyruvate (PEP), the substrate of CO_2_-fixation reaction via PEP carboxylase, seem to reduce under Lys-treatment in the earlier phases (**Table 1**).

Overall, the lowest average foldchange was measured for dihydropteroate (0.14; **Table 1**), an intermediate of folate biosynthesis. Folate derivatives are required for biosynthesis of D-pantothenic acid (CoA), which had the third highest average foldchange of 13.02 (**Table 1**), implying a drain of dihydropteroate toward folate derivatives to produce D-pantothenic acid (CoA). Furthermore, folate derivatives contribute to amino acid metabolism via the methionine cycle and L-glycine/L-serine (alanine) biosynthesis – all of which over-accumulated – and to nucleotide metabolism (Mills *et al*., 2020). Interestingly, several metabolites of nucleotide metabolism accumulated as well (**Table 1; Suppl. Fig. S4**), especially those of lower energy (guanosine, adenosine, cytidine, guanine, adenine, CMP, AMP, and GMP), suggesting an energy crisis under Lys-treatment. This is in agreement with the drop of the photosynthetic activity of *Synechocystis* after Lys-treatment (**Fig. 1B**).

Additionally, we observed an overaccumulation of oxidized glutathione among the highest foldchange of 7.00 (**Table 1**), which is known as a sign of oxidative stress. To confirm this observation, we quantified the production of reactive oxygen species (ROS) after 24 h, 48 h, 72 h, 96 h, and 168 h of Lys-treatment using the H2DCFDA fluorescent marker as previously described (Haffner *et al*., 2023). The first 24 h were comparable for both conditions, however, between 48-96 h the fluorescence was 100-200% times higher for Lys-treated cells (**Fig. 1G**), indicating more ROS and by extension higher oxidative stress.

### 2.3 Escaping L-lysine toxicity

Importantly, the growth of *Synechocystis* cells was only arrested for roughly 72 h when grown with 25 µM of Lys and resumed after (**Fig. 1A**), suggesting that the cells acquire Lys-resistance. To identify which genes might contribute to *Synechocystis* cells escaping the Lys toxicity, we grew a pooled library of CRISPRi *Synechocystis* mutants either in the presence or absence of 25 µM Lys. In this library, up to five sgRNAs target each annotated gene of *Synechocystis* by CRISPR inhibition (Miao & Jahn *et al*., 2023). NGS libraries were prepared from samples before and after addition of aTc and Lys and sequenced to track abundances of library members over time. Based on the abundance changes, the fitness of the genes targeted by the respective set of sgRNAs was calculated, as described previously (Miao & Jahn *et al*., 2023). Correlation between sgRNA counts of different CRISPRi samples showed that Lys-treatment had a stronger impact on library composition than the control treatment (**Suppl. Fig. S5**). As observed for WT cultures, the pooled library showed a transient growth arrest after addition of Lys (**Suppl. Fig. S6**). In both growth regimes, with and without Lys, known essential genes were depleted over time, e.g., genes encoding ribosomal proteins (**Suppl. Tables S3 and S4**). This was verified by GSEA on basis of gene ontology and KEGG terms (**Supp. Tables S5 to S10**). This confirms the suitability of the CRISPRi approach to identify genes which are important for a given growth condition. The sgRNAs targeting genes associated with several KEGG pathways related to amino acid biosynthesis and translation were enriched in the control library compared to the Lys-treated library (**Suppl. Table S10, Supp. Fig. 7**). An enrichment of these sgRNAs in the control condition relative to the Lys-Lys-treated cells than the control cells.

Suppression of several genes led to a relative fitness advantage in the Lys-treated library compared to the control library (**Fig. 3A**, **Table 2**). We identified the set of genes with the strongest difference by subtracting the fitness values for the two conditions (**Table 2**). Genes with a high negative fitness difference show the strongest fitness advantage after Lys-treatment compared to the control condition. This set of genes included *bgtA* (*slr1735*) and *bgtB* (*sll1270*), which encode the basic amino acid and glutamine transporter, Bgt-system (**Fig. 3A**). *Synechocystis bgtA* and *bgtB* deletion mutants are known to survive toxic concentrations of L-glutamine (Quintero *et al*., 2001). Other genes that specifically gave fitness in the Lys-treated library involved *uirR* (*slr1213*), which encodes the DNA-binding response regulator UirR (Song *et al*. 2011), and *pirA* (*ssr0692*), a regulatory protein which regulates nitrogen flow towards the OA cycle (Bolay *et al*., 2021). Many of the genes which showed different fitness patterns between both growth conditions involved hypothetical and unknown proteins, e.g., *sll1200*, *slr0262*, and *slr5017*. Also, several genes showed strong negative fitness under Lys-treatment (**Table 2**), e.g., the twitching motility protein PilT2 (*sll1533*). To validate our library screen, we compared the growth, photosynthetic activity, and pigmentation of a *bgtA* mutant (Δ*bgtA*) to WT *Synechocystis* with and without 25 µM Lys. In contrast to WT cells, the Δ*bgtA* mutant grew comparably under both conditions and showed similar photosynthetic activity and pigmentation (**Fig. 3B and 3C, Suppl. Fig. S8**), supporting that Bgt-system is a main Lys importer and indicating that Lys has to be taken up to elicit a toxic response and further validating our CRISPRi library screen.

**Figure 3:**
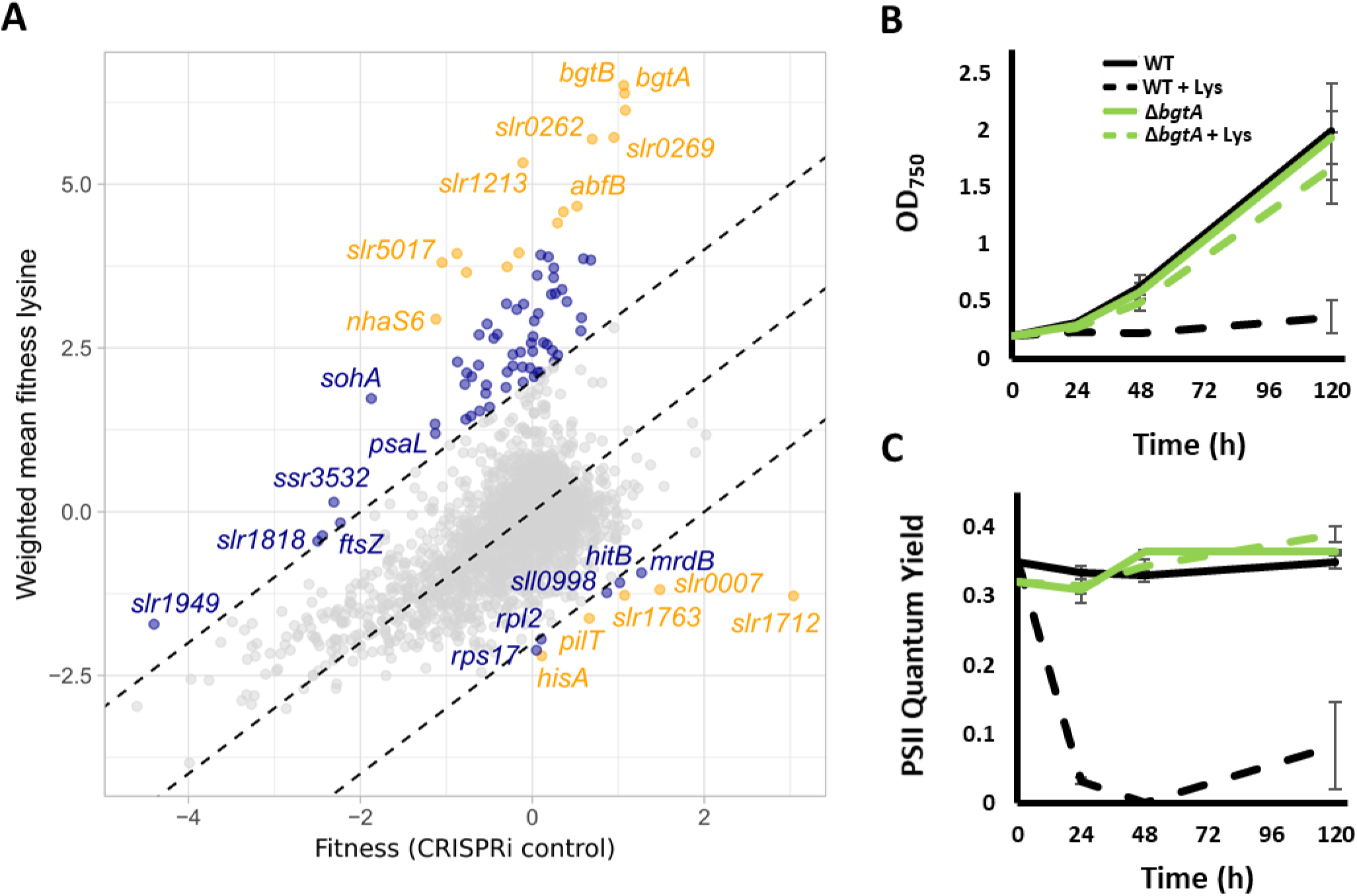
Pooled CRISPRi library screen with and without 25 µM Lys. **A:** Dot plot fitness with Lys plotted versus fitness without Lys of CRISPRi knockdown of all protein-coding genes. **B & C:** validation of CRISPRi screen for Δ*bgtA.* **B:** Growth represented by the mean of OD_750_ ± SD from biological triplicates. **C:** Photosynthetic efficiency measured by PAM fluorometer showing the PSII quantum yield ± SD from biological triplicates.

**Table 2:** Overview of genes which showed the highest difference between the control cultivation and the cultivation with added Lys in the CRISPRi library. Genes with a fitness difference of below −4 and the five genes with the highest values are given. The complete list of genes is given in Supplementary Tables (S3 and S4). A negative fitness difference implies that sgRNAs targeting a specific gene were more enriched in the Lys-treated library than in the control library, corresponding to a stronger positive fitness effect after Lys addition than in the control condition. A positive fitness difference implies the opposite.

| Locus tag | Gene name | Gene product | Fitness (control) | Fitness (25 μM Lys) | Fitness difference (control – 25 μM Lys) |
| --- | --- | --- | --- | --- | --- |
| *sll1270* | *bgtB* | periplasmic substrate-binding and integral membrane protein of the ABC-type Bgt permease for basic amino acids and glutamine BgtB | 1.06 | 6.51 | -5.44 |
| *slr1213* | *uirR* | two-component response regulator AraC subfamily | -0.11 | 5.33 | -5.44 |
| *slr1735* | *bgtA* | ATP-binding subunit of the ABC-type Bgt permease for basic amino acids and glutamine | 1.07 | 6.39 | -5.31 |
| *sll1200* | *sll1200* | hypothetical protein | 1.08 | 6.13 | -5.05 |
| *slr0262* | *slr0262* | unknown protein | 0.70 | 5.69 | -4.99 |
| *slr5017* | *slr5017* | hypothetical protein | -1.05 | 3.80 | -4.85 |
| *sll0062* | *sll0062* | hypothetical protein | -0.88 | 3.94 | -4.82 |
| *slr0269* | *slr0269* | hypothetical protein | 0.95 | 5.71 | -4.76 |
| *ssl1807* | *ssl1807* | hypothetical protein | -0.77 | 3.65 | -4.42 |
| *slr1935* | *slr1935* | hypothetical protein | 0.36 | 4.58 | -4.22 |
| *slr0518* | *abfB* | similar to alpha-L-arabinofuranosidase B | 0.52 | 4.66 | -4.14 |
| *slr0579* | *slr0579* | unknown protein | 0.29 | 4.41 | -4.11 |
| *ssr0692* | *pirA* | P<sub>II</sub>-interacting regulator of arginine synthesis (PirA) | -0.16 | 3.95 | -4.11 |
| *sll0556* | *nhaS6* | Na<sup>+</sup>/H<sup>+</sup> antiporter | -1.12 | 2.94 | -4.06 |
| *slr0096* | *slr0096* | low affinity sulfate transporter | -0.29 | 3.74 | -4.03 |
| *sll1533* | *pilT2* | twitching motility protein | 0.66 | -1.63 | 2.29 |
| *slr0652* | *hisA* | phosphorybosilformimino-5-amino-phosphorybosil -4-imidazolecarboxamideisomerase | 0.11 | -2.20 | 2.31 |
| *slr1763* | *slr1763* | probable methyltransferase | 1.07 | -1.28 | 2.35 |
| *slr0007* | *slr0007* | probable sugar-phosphate nucleotidyltransferase | 1.48 | -1.19 | 2.67 |
| *slr1712* | *slr1712* | hypothetical protein with homology to DNA endonuclease RpnA | 3.04 | -1.28 | 4.32 |

### 2.4 Potential involvement of second messenger c-di-AMP in cell wall homeostasis

The *dacA* gene (*sll0505*), encoding the only di-adenylate cyclase that synthesizes the second messenger c-di-AMP (cyclic dimeric adenosine monophosphate), is transcribed from an operon with two genes related to peptidoglycan synthesis; *lysA* (*sll0504*) and *uppS* (*sll0506*) (Agostoni *et al*., 2018; Commichau *et al*., 2019; Selim *et al*. 2021). The genetic locus of *dacA* suggests a possible link of c-di-AMP to cell wall homeostasis and by extension Lys metabolism. Furthermore, the CRISPRi analysis revealed that the sgRNA of the Na^+^/H^+^ antiporter *nhaS6* (encoded by *sll0556*), which is strongly upregulated in the knockout mutant Δ*dacA*, was among the most enriched sgRNA in Lys-treatment. Oppositely, the sgRNA of the twitching motility protein *pilT2* (encoded by *sll1533*), which is strongly downregulated in Δ*dacA*, was among the most depleted sgRNA in Lys-treatment (**Table 2**) (Mantovani *et al*., negatively influence on the cell fitness under Lys stress.

To investigate the relationship between c-di-AMP and Lys toxicity, Δ*dacA* growth was compared to WT growth on a drop dilution assay in BG11^N^ with and without 20 µM Lys, a concentration that does not influence on WT negatively. In addition, the complementation strain Δ*dacA*::P*petE*-*dacA* and the c-di-AMP overexpression strain WT::P*petE*-*dacA* (Samir *et al*. 2023), both of which express *dacA* under the copper-inducible promoter P*petE* (Englund *et al*., 2016; Selim *et al*. 2021), were tested. Under both conditions, WT and WT::P*petE*-*dacA* grew comparably, implying that high c-di-AMP does not influence on Lys-toxicity under tested Lys concentration. In contrast, no visible growth was observed for Δ*dacA* with Lys, while Δ*dacA*::P*petE*-*dacA* strain was able to partially complement the Δ*dacA* growth defect (**Fig. 4**), supporting a role for c-di-AMP in the overall cell fitness and regulating cell wall integrity.

**Figure 4:**
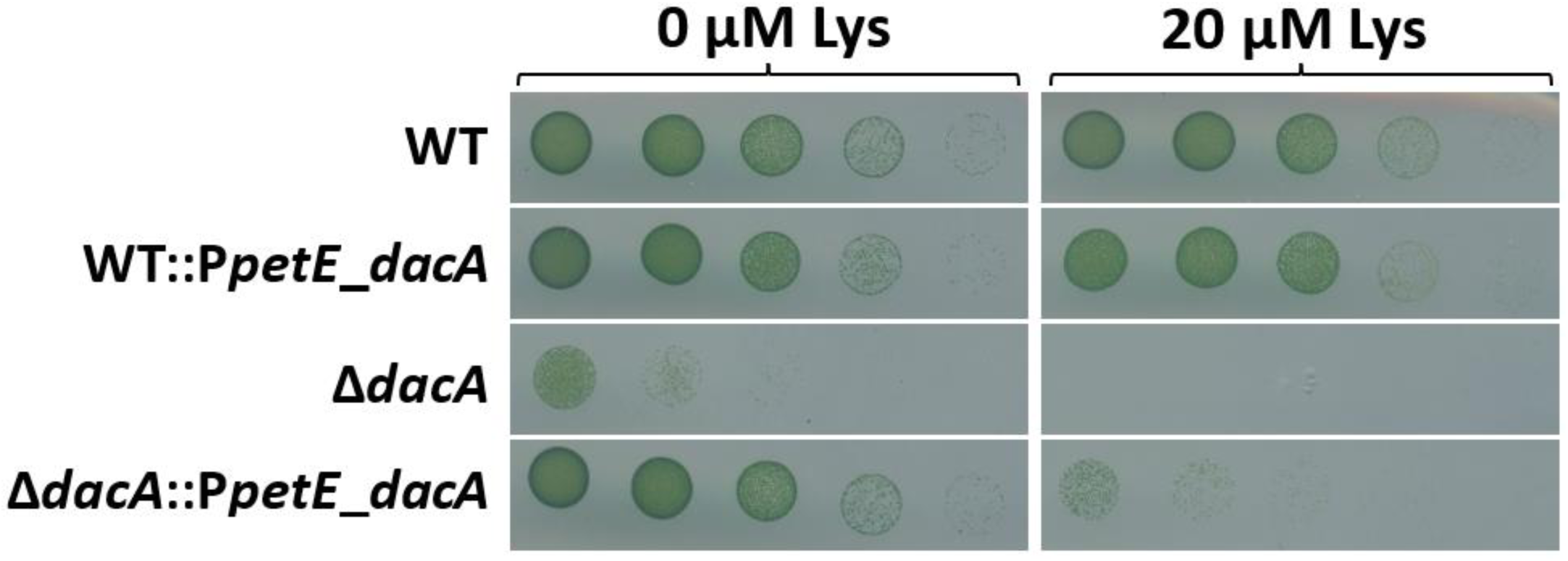
Phenotypic characterization of c-di-AMP free mutant Δ*dacA* under Lys toxicity. Drop assay of WT, WT::P*petE*-*dacA*, Δ*dacA*, and Δ*dacA*::P*petE*-*dacA* in BG11^N^ with and without 20 µM Lys after 168 h (1 week). Cells were dropped in serial 1:10 dilutions starting from OD_750_ 1 (left).

## 3. Discussion

Building upon previous research demonstrating the detrimental effects of Lys incorporation on cell wall integrity and subsequent cell death in *M. aeruginosa* (Kim *et al*., 2023; Yan *et al*. 2023), we investigated the early metabolic response of the model cyanobacterium *Synechocystis* sp. PCC 6803 to Lys exposure. Our study provides physiological and metabolic evidence of the same phenomenon as occurring in *M. aeruginosa*. Additionally, we show other negative physiological influences of Lys-toxicity including cell enlargement, oxidative stress, and damage to the photosystems, which all contribute to growth arrest. In the metabolism, high levels of the peptide chain constituents L-/D-alanine, mDAP, and D-Ala-D-Ala are indicative of peptidoglycan disruption. The observed overaccumulation of mDAP provides further support for the absence of negative feedback regulation on Lys synthesis, and corroborates previous findings showing transcriptional upregulation of key enzymes, namely LysC, DapF, and LysA, involved in the mDAP pathway following Lys exposure in *M. aeruginosa* (Kim *et al*., 2023). Moreover, our results align with previous observations indicating that Lys does not impede D-Ala-D-Ala ligase activity (Kim *et al*., 2023), as evidenced by the overaccumulation of D-Ala-D-Ala in our metabolomic analysis. Lys also provoked alterations in amino acid levels upstream of the alanine pathway and other amino acid pathways, notably elevated levels of tyrosine, tryptophan, valine, and iso-/leucine, implying a broader influence on amino acid metabolism. The broader influence of Lys on amino acid metabolism could be, *e.g*., through Lys riboswitches, which are widespread in bacteria and are known to influence the levels of multiple amino acids (Serganov & Patel, 2009). Although Yan *et al*. (2023) reported high levels of OA cycle metabolites in *M. aeruginosa* after 72 h of Lys exposure, these metabolites were not as affected in *Synechocystis* as other metabolites, but showed a clear trend of upregulation after Lys-treatment. Remarkably, an apparent drop of OA was observed the first 24 h for OA cycle and GS-GOGAT cycle metabolites. This could be due to the photo-inhibitory effect of Lys, which could potentially downregulate the GS-GOGAT cycle and consequently the nitrogen flow towards L-arginine biosynthesis (Reyes *et al*., 1995; Cassier-Chauvat *et al*., 2014; Forchhammer & Selim, 2020). Nevertheless, these metabolites were more abundant in Lys-treated cells, which supports findings that the OA cycle serves as a buffer for nitrogen oscillations (Zhang *et al*., 2018). This is also in accordance with our CRISPRi library screen, showing that knocking-down *pirA* increases the cellular fitness under Lys-treatment. PirA is known to regulate the flux into the OA cycle by sequestering PII signaling protein from its primary target, N-acetyl-L-glutamate kinase (NAGK; key enzyme of arginine biosynthesis), thereby tuning the nitrogen flow towards the OA cycle (Forchhammer *et al*. 2022). Thus, Δ*pirA* mutant shows more nitrogen flow towards the OA cycle with accumulation of arginine, ornithine, and citrulline metabolites (Bolay *et al*., 2021).

Interestingly, out of all the metabolites measured, Lys itself was one of the most abundant metabolites. More importantly, its levels were noticeably high at all measured time points, peaking at 48 h and slowly decreasing thereafter. This indicates slow Lys catabolism and raises the question whether *Synechocystis* possesses any dedicated Lys catabolic pathways. In addition to amino acid metabolism, our findings suggest a potential connection between Lys integration and nucleotide metabolism, possibly through the methionine and folate cycle. Specifically, we observed highly elevated levels of SAM and notably low levels of the folate precursor dihydropteroate; the only metabolite strongly downregulated in response to Lys-treatment. This observation hints at intricate metabolic crosstalk between amino acid metabolism and nucleotide biosynthesis pathways, implicating potential regulatory nodes and metabolic flux redirection.

Complementing our metabolomic analysis, we employed a CRISPRi-based approach to pinpoint genes contributing to the resistance of Lys toxicity in *Synechocystis*. The subunits of the Bgt permease for basic amino acids and glutamine were among the top hits, underscoring the necessity of Lys uptake for eliciting its toxic response. Our growth experiments confirmed that a BgtA mutant could grow normally on Lys. Another top hit was the DNA-binding response regulator UirR, which is inhibited by ethylene (Kuchmina *et al*., 2017) or activated in response to UV-A light. Upon activation, UirR induces the expression of the response regulator LsiR, associated with negative UV-phototaxis (Song *et al*., 2011), and of the small regulatory RNA CsiR1, which exists as a discrete transcript and is implicated in carbon acclimation response (Klähn *et al*., 2015). Additionally, our screen identified two genes likely regulated by the second messenger c-di-AMP; *pilT2* and *nhaS6* (Mantovani *et al*., 2022; Samir *et al*. 2023). Of particular interest is the diadenylate cyclase gene (*dacA*) responsible for producing c-di-AMP, located in the same operon as *uppS* and *lysA*, which are crucial for peptidoglycan synthesis. Intriguingly, in *Bacillus subtilis*, the synthesis and degradation of c-di-AMP influences the resistance to ß-lactams; a class of antibiotics that inhibits peptidoglycan crosslinking (Luo & Helmann, 2011). Similarly, our knockout analysis revealed that Δ*dacA* displayed heightened sensitivity to Lys compared to WT. This increased sensitivity may arise from a direct role of c-di-AMP in cell wall homeostasis; however, given the cellular enlargement induced by Lys exposure and the conserved role of c-di-AMP in osmoregulation (Agostoni *et al*., 2018; Selim *et al*. 2021; Mantovani *et al*. 2023), the absence of c-di-AMP might have exacerbated osmotic stress instead. The precise link between c-di-AMP and cell wall homeostasis in *Synechocystis* remains elusive and warrants further investigation. Additionally, a previously link between c-di-AMP and ROS, which accumulates in the cell under Lys-treatment (**Fig. 2**), was observed (Haffner *et al.,* 2023), further supporting a role for c-di-AMP in regulating Lys toxicity.

Overall, our study reveals the early metabolic and physiological responses of *Synechocystis* sp. PCC 6803 to L-lysine exposure, mirroring detrimental effects observed in *M. aeruginosa*. Metabolomic analysis highlights disruptions in peptidoglycan biosynthesis, supported by physiological evidence of cell enlargement, oxidative stress, and damage to photosystems. Genetic analysis identifies key genes involved in the cellular response to Lys toxicity, providing insights into potential regulatory pathways and avenues for future research in controlling cyanobacterial blooms.

## 4. Materials and Methods

### Cultivation of *Synechocystis* sp. PCC 6803

*Synechocystis* strains (**Table S1**) were grown in BG11^N^ medium supplemented with NaNO_3_ as nitrogen source (**Table S2**) as described previously (Selim *et al*., 2018, 2021, 2023). For nitrogen-limiting conditions, NaNO_3_ was removed (BG11^0^). Solid medium contained 1% (w/v) Bacto^TM^ agar. The standard cultivation consisted in growth at 28 °C with constant illumination of 40-60 µE and constant shaking at 120 rpm. To prevent light stress, cultures were grown at low light (20 µE) the first 24 h. Pre-cultures of the Δ*bgtA* (*slr1735*) mutant were kept under selection pressure with the appropriate antibiotics (**Table S1**). The Δ*dacA*, Δ*dacA*::P*petE*-*dacA* and WT::P*petE*-*dacA* mutants were created as described previously (Selim *et al*. 2021; Samir *et al*. 2023). The primers and plasmids used are listed (**Table S1**).

### Growth assays

Pre- and main cultures were inoculated at an OD_750_ of 0.2 (early exponential growth phase). Pre-cultures were harvested for main cultures at an OD_750_ of ∼0.8 (late exponential growth phase) by centrifugation at 4000 g for 10 min and washed with the appropriate BG11^N^ medium. Growth assays included cultivation in 24-well-plates (2 ml cultures) and cultivation in flasks (50 ml and 250 ml cultures). For flask cultures, biological triplicates were used; for well-plates, biological duplicates were used. Growth was assessed with and without Lys at micromolar concentrations by measuring OD_750_ with an Ultrospec III Spectrophotometer (Pharmacia LKB; New Jersey, USA).

### Whole-cell spectrum

To determine the degree of pigmentation (*i.e*., the integrity of the photosystems), the whole cell absorbance spectrum (WCS) was measured from 320 - 750 nm using the Specord 205 (Analytic Jena AG; Jena, Germany) as described previously (Selim *et al*. 2018, 2021, 2023). Samples were diluted to an OD_750_ between 0.1 – 0.5 for accurate measurement. The spectra were normalized to OD_750_ by dividing values with the 750 nm absorbance value.

### Pulse-Amplitude-Modulation Fluorometry

The photosystem II (PSII) activity was measured with a WATER-PAM chlorophyll fluorometer (Walz GmbH; Effeltrich, Germany), as described previously (Selim *et al*. 2018). Samples were diluted in about 2 ml (no visible coloration) and normalized to a minimal base fluorescence between 400-500. The first measurement was omitted, and the next three were treated as technical triplicates.

### Microscopy

*Synechocystis* cells were stained with 10 μM DiBAC_4_(3) (Bis-(1,3-Dibutylbarbituric Acid)- trimethine oxonol) (AAT Bioquest (Hamburg, Germany; cat. no. 21411; 10 mM stock dissolved in DMSO, diluted in MilliQ to 100 µM) for 30 min in the dark. 5 μL of stained cells were dropped on an agarose-coated microscopy slide (1 % (w/v) in H_2_O). Imaging was performed with the Zeiss Axio Imager M2 (Oberkochen, Germany) with an 100x /1.3 oil objective. A green fluorescent protein (GFP) filter (excitation peak: 488 nm; emission peak 510 nm) was used to detect DiBAC_4_(3).

### Reactive oxygen species (ROS) quantification

ROS were quantified using the H2DCFDA fluorescent marker as previously described (Diamond *et al*. 2017; Haffner *et al*. 2023). *Synechocystis* cells were normalized to OD_750_ 0.4 and H2DCFDA was added to a final concentration of 5 µM and incubated for 1 h at 30 °C in the dark. 200 µL of each sample were transferred into a 96-well plate and fluorescence was measured (480 nm excitation; 520 nm emission) using the CLARIOstar® plate reader (BMG Labtech; Ortenberg, Germany).

### CRISPR interference (CRISPRi)

A *Synechocystis* CRISPRi library (Miao & Jahn *et al*., 2023), in which up to five sgRNAs target each annotated gene, was used to measure fitness effects of a transcriptional knockdown of the respective genes under different growth conditions. To this end, the pooled library was cultivated in 8-tube Multi-Cultivator MC-1000-OD bioreactors (Photon System Instruments, Drasov, CZ) in turbidostat mode as described previously (Miao & Jahn et al., 2023). Light intensity was set to 45 µE and ambient air was used as gas feed. Cultures were grown at 30°C in BG-11 medium (Rippka et al., 1979) substituted with 20 mM HEPES (pH 7.8), 25 µg mL^-1^ spectinomycin and 50 µg mL^-1^ kanamycin. The turbidity threshold was set to OD_720nm_ = 0.2. Cultures were diluted by addition of fresh medium if this threshold was exceeded for two measurements in a row. Expression of dCas9 and sgRNAs was induced by addition of 0.5 µg mL^-1^ anhydrotetracycline (aTc). Samples corresponding to generation 0, *i.e*., the point of reference for further comparisons, were collected 24 h after aTc addition and before adding 25 µM Lys to two of the four replicate cultivations. At the 8th generation, cells were harvested for the control cultivation without added lysine. For the cultivation with added lysine, cells were harvested at the same time point. Cells were harvested, samples processed and the next generation sequencing (NGS) library was prepared and sequenced as described previously (Miao & Jahn et al., 2023). Data analysis was performed using the nf-core-crispriscreen pipeline (https://github.com/MPUSP/nf-core-crispriscreen, commit: c7c6563eb269bc3426e7481a8324788bc1006306). Further code for data processing and analysis is available on GitHub (https://github.com/ute-hoffmann/Lys_Synechocystis) and Zenodo (doi: 10.5281/zenodo.11093669). Gene set enrichment analysis (GSEA) was performed using clusterProfiler (v4.8.3) (Wu *et al*., 2021; Yu *et al*., 2012). Fitness data for all mutants can be accessed on an interactive web application (https://m-jahn.shinyapps.io/ShinyLib/).

### Metabolomics Sampling

For metabolome investigation of *Synechocystis* growing with and without 25 µM Lys (BG11^N^), samples were collected before addition of Lys and 24 h, 48 h, 72 h, and 120 h after. To ensure enough cell material was available, cells were cultured in 250 ml. Cells (OD_750_ of 6.0) were collected on glass microfiber membrane filters (Ø 25 mm, pore size 1.2 µm, GE Healthcare Life Sciences) via vacuum filtration and washed with 10 ml BG11^N^. The filters were immediately transferred into 2 ml tubes, frozen in liquid nitrogen, and stored at −80 °C for future extraction. Culture volumes were maintained equally by subtracting differences in extraction volume. To extract the metabolites for LC-MS/MS analysis, 1 ml of ice-cold quenching solution (acetonitrile:methanol:water; 2:2:1) was distributed into glass vials and the frozen filters were submerged completely. The vials were then incubated at −20 °C for 30 min and the contents were subsequently mixed by pipetting. 500 µl were transferred into a 1.5 ml tube on ice and centrifuged (14 000 g, 4 °C, 15 min). 400 µl of the supernatant were transferred into a new 1.5 ml tube and stored at −80 °C until analysis. Targeted LC-MS/MS analysis was performed with a triple quadrupole mass spectrometer as described previously (Guder *et al*. 2017). Raw measurement data can be found in (**Suppl. Table S11**).

## Data availability

Raw sequencing data were deposited at the European Nucleotide Archive (ENA accession number PRJEB75388). Code used for data processing and analysis are available online in GitHub and Zenodo repositories (https://github.com/ute-hoffmann/Lys_Synechocystis and doi: 10.5281/zenodo.11093669). Fitness data for all mutants can be accessed on an interactive web application (https://m-jahn.shinyapps.io/ShinyLib/).

## Author contributions

AA and KAS conceived and initiated the research; KAS supervised the whole study and designed the experiments with AA; AA performed most of the physiological and biochemical experiments, while RM and UAH performed CRISPRi library screen and JR ran the LC-MS/MS metabolome analysis; PH provided and supervised the CRISPRi screen, while HL supervised the metabolome analysis; AA and KAS evaluated and interpreted the results; AA prepared the figures and wrote the first draft of the manuscript with inputs from KAS. All authors approved the final version of the manuscript.

## Acknowledgements

The project was funded by the German Research Foundation (DFG) as part of the priority research program SPP2389 (SE 3449/1-1) to KAS. KAS and AA gratefully acknowledges Karl Forchhammer for continued support during and after this project, and acknowledges the infrastructural support by the Cluster of Excellence “Controlling Microbes to Fight Infections (CMFI)” (EXC2124–390838134), the CRC 1535 (MibiNet), and the Excellence Strategy of the German Federal and Baden-Württemberg State Governments to KAS (Projektförderung: PRO-SELIM-2022-14). RM, UAH, and PH acknowledge funding from the Novo Nordisk Foundation (NNF20OC0061469).

---

**Suppl. Fig. S1**

**Supplementary Figure S1.**
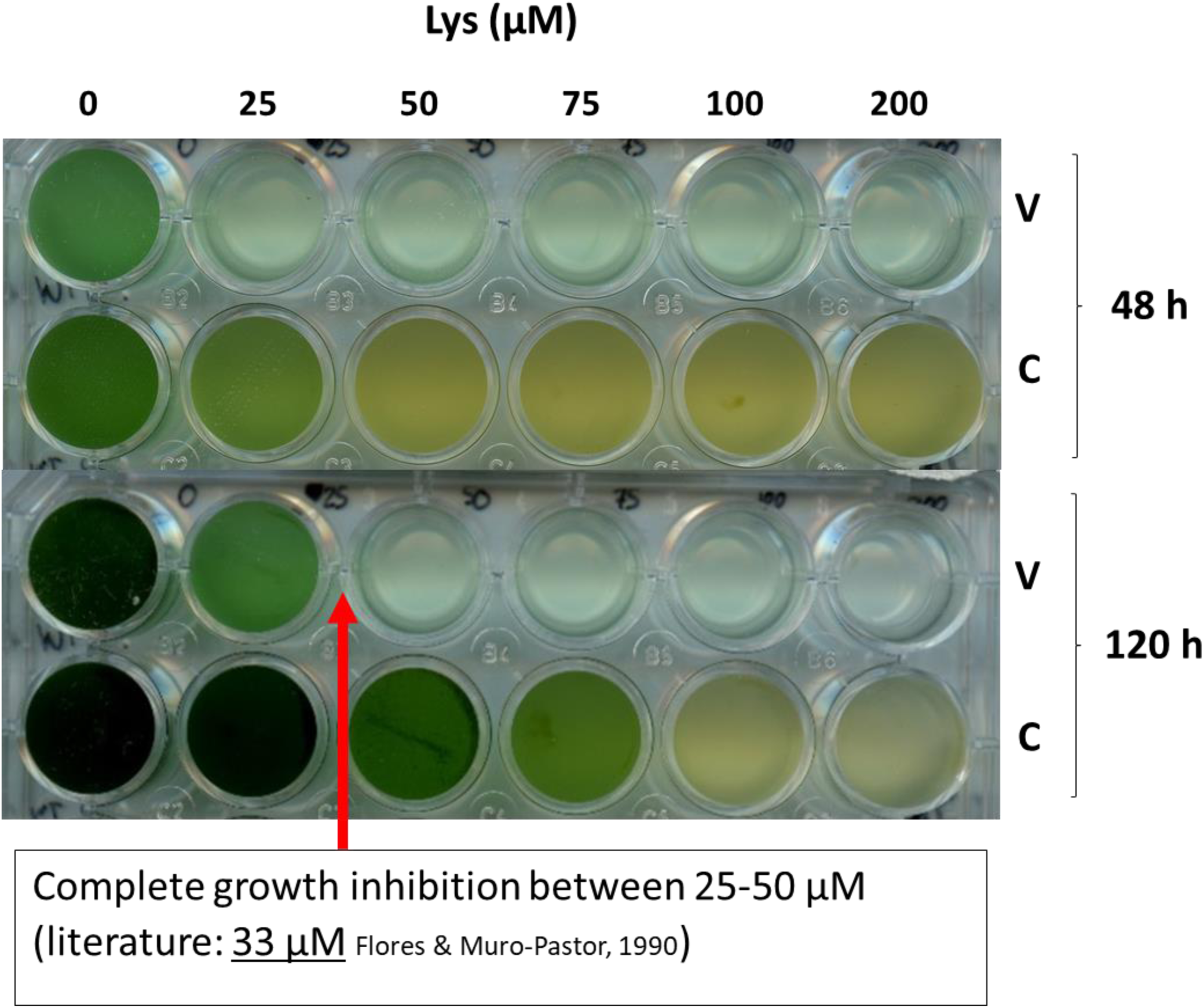
Lysine titration (0-200 µM) of *Synechocystis* WT cells in vegetative and chlorotic state. The vegetative cells (V) were inoculated in BG11^N^ with different concentrations of Lys as indicated. The chlorotic cells (C) were resuscitated with the same concentrations of Lys in BG11^N^ media.

**Suppl. Fig. S2**

**Supplementary Figure S2.**
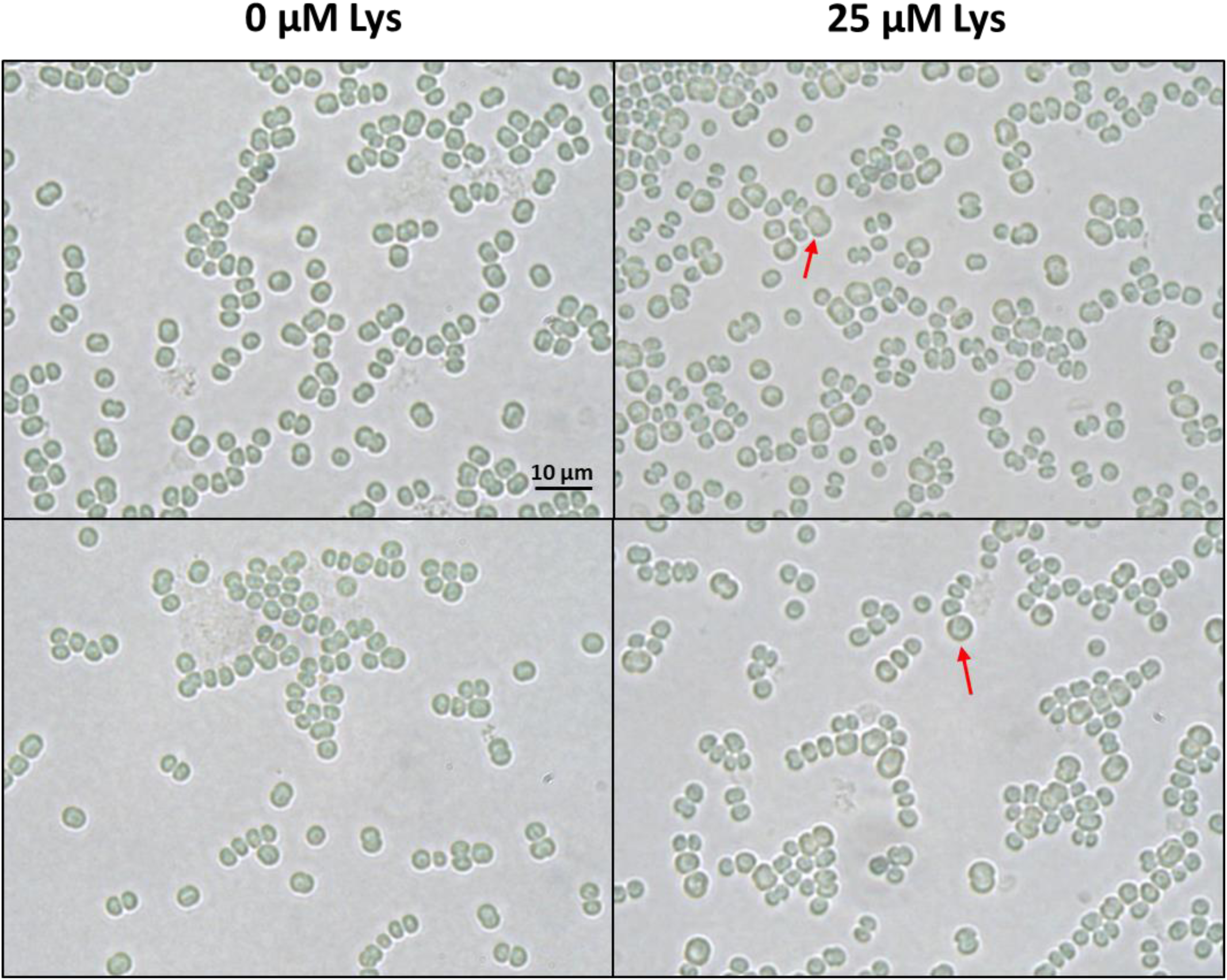
Lysine titration (0-200 µM) of *Synechocystis* WT cells in vegetative and chlorotic state. The vegetative cells (V) were inoculated in BG11^N^ with different concentrations of Lys as indicated. The chlorotic cells (C) were resuscitated with the same concentrations of Lys in BG11^N^ media.

**Suppl. Fig. S3**

**Supplementary Figure S3:**
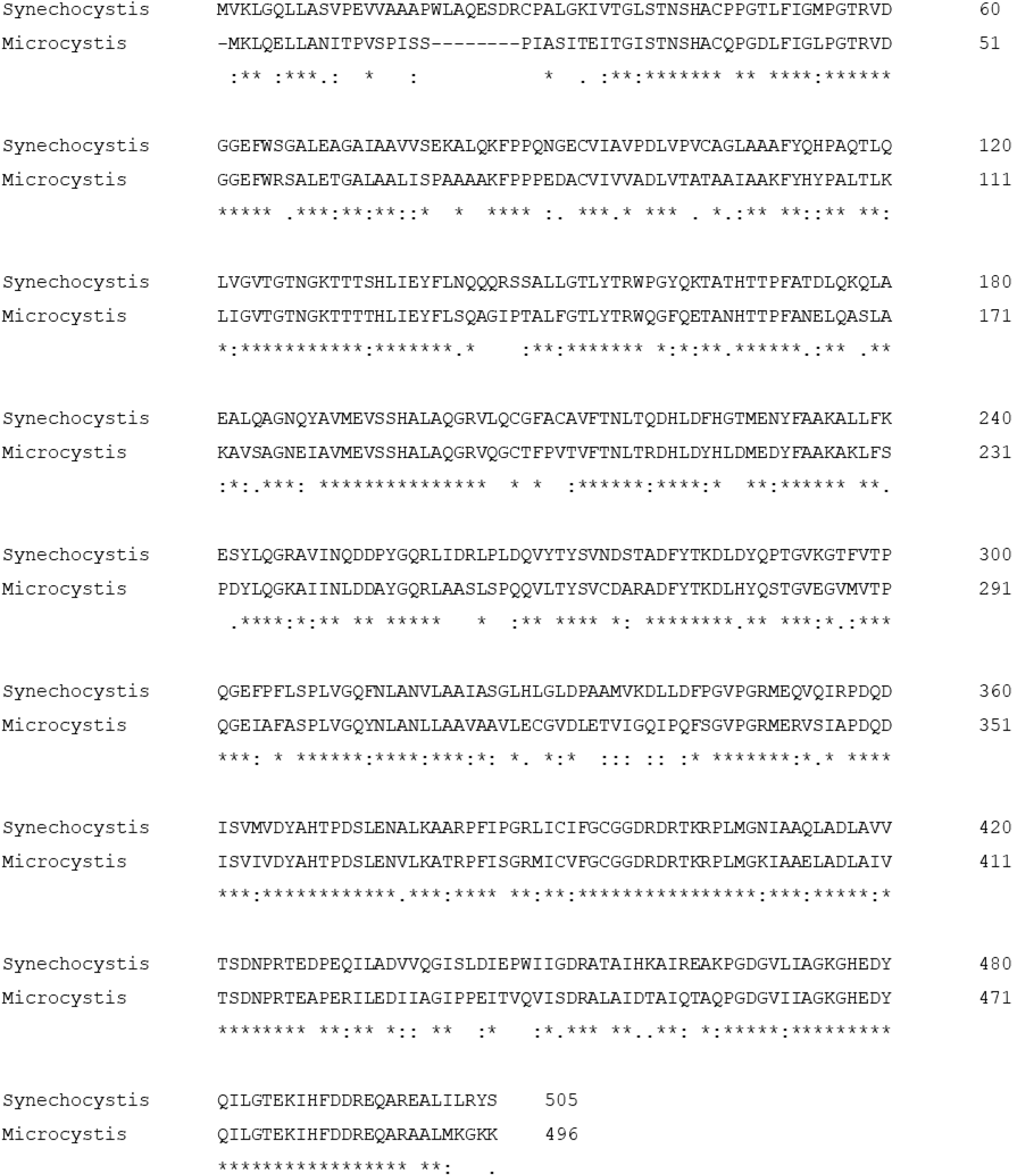
Multiple sequence alignment generated by Clustal Omega for MurE from *Synechocystis* sp. PCC 6803 and *Microcystis aeruginosa* PCC 7806, showing 67% sequence identity.

**Suppl. Fig. S4**

**Supplementary Figure S4:**
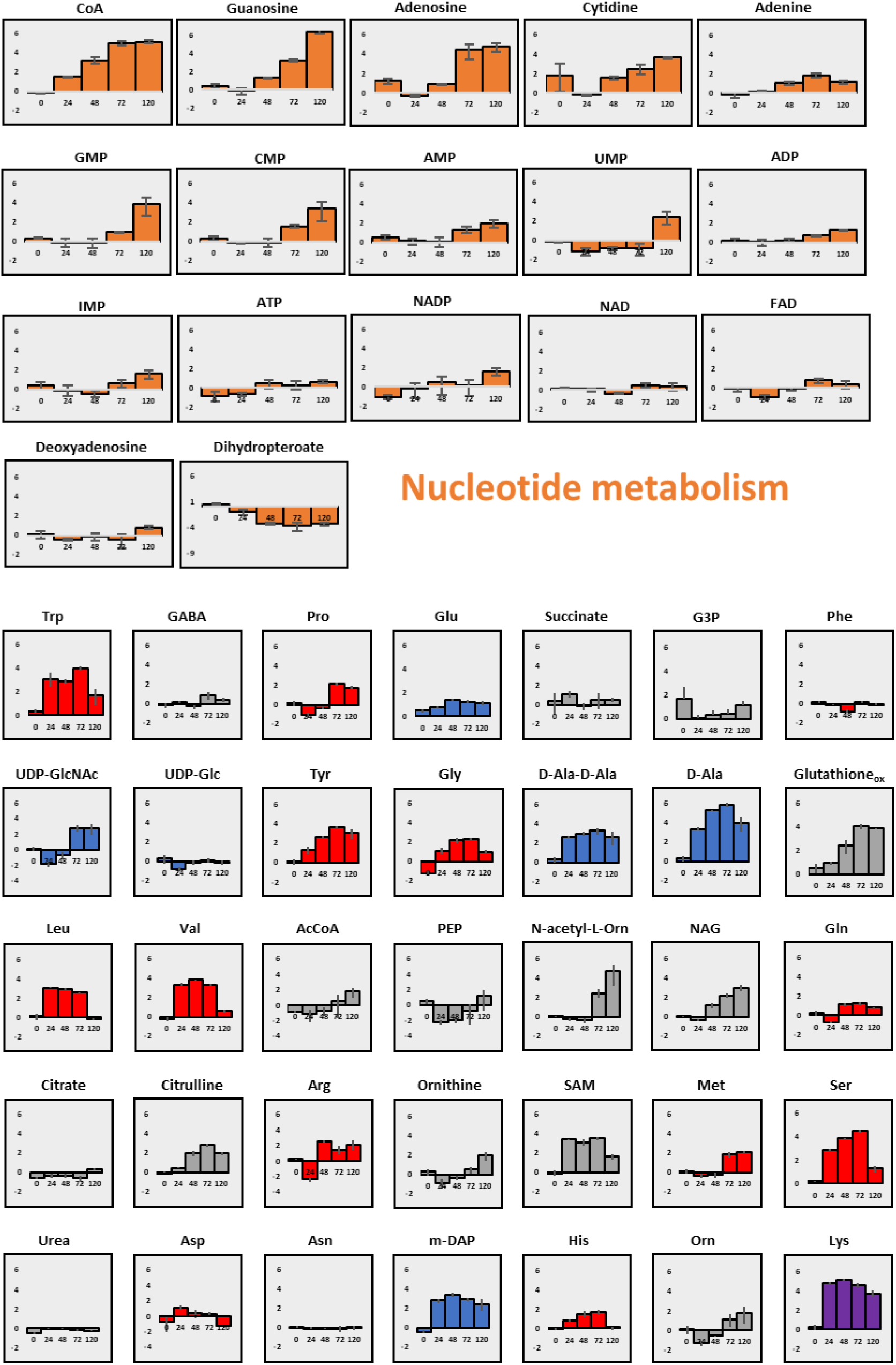
Collection of the graphs from Figure 2 and for metabolites related to nucleotide metabolism. X-axis: Time (h). Y-axis: log_2_(foldchange). Nucleotide metabolism in **orange**, Lys in **purple**, cell wall metabolism in **blue**, proteinogenic amino acids in **red**, and other measured metabolites in **grey**.

**Suppl. Fig. S5**

**Supplementary Figure S5:**
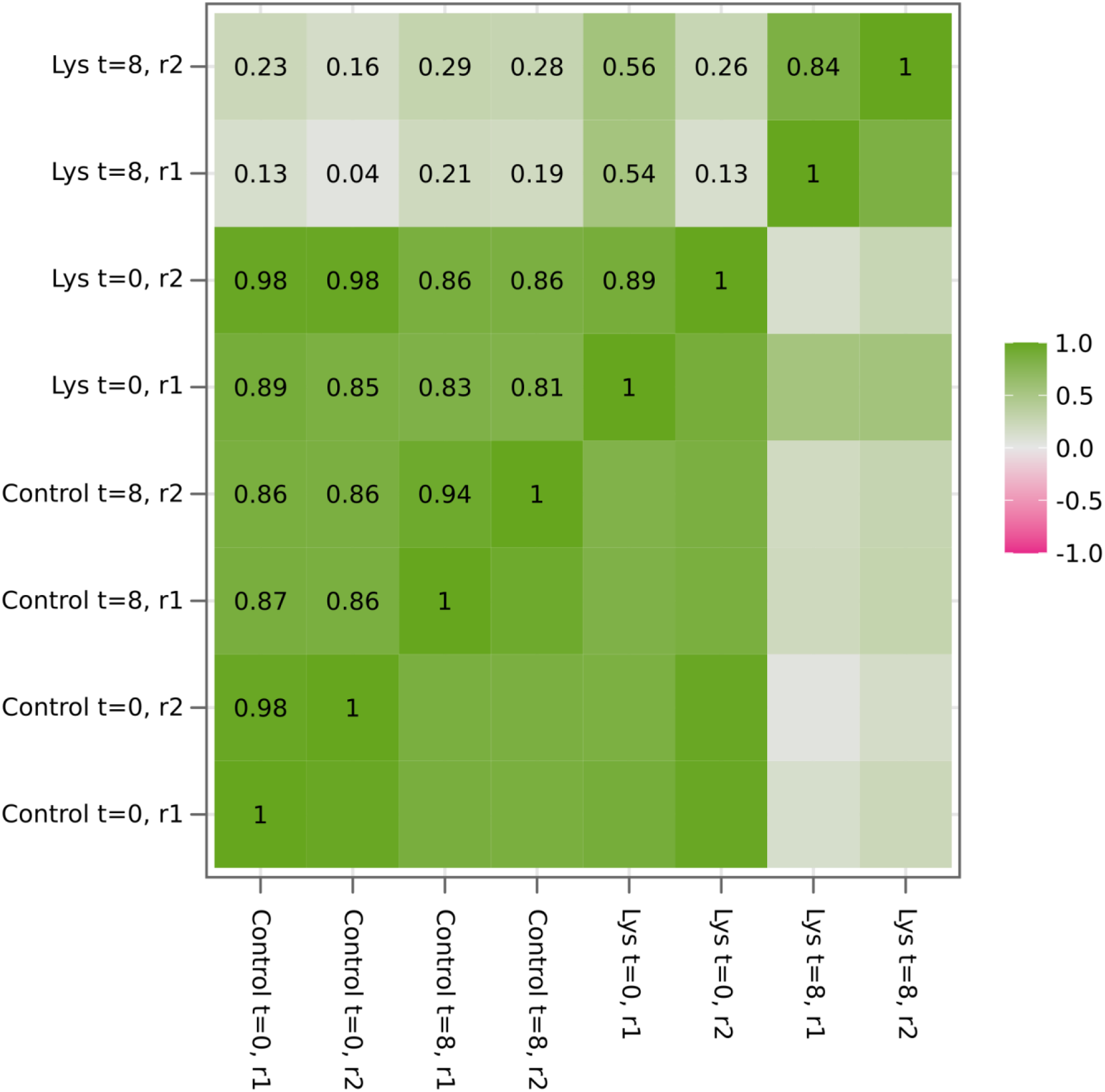
Pearson correlations between sgRNA counts of different CRISPRi samples.

**Suppl. Fig. S6**

**Supplementary Figure S6:**
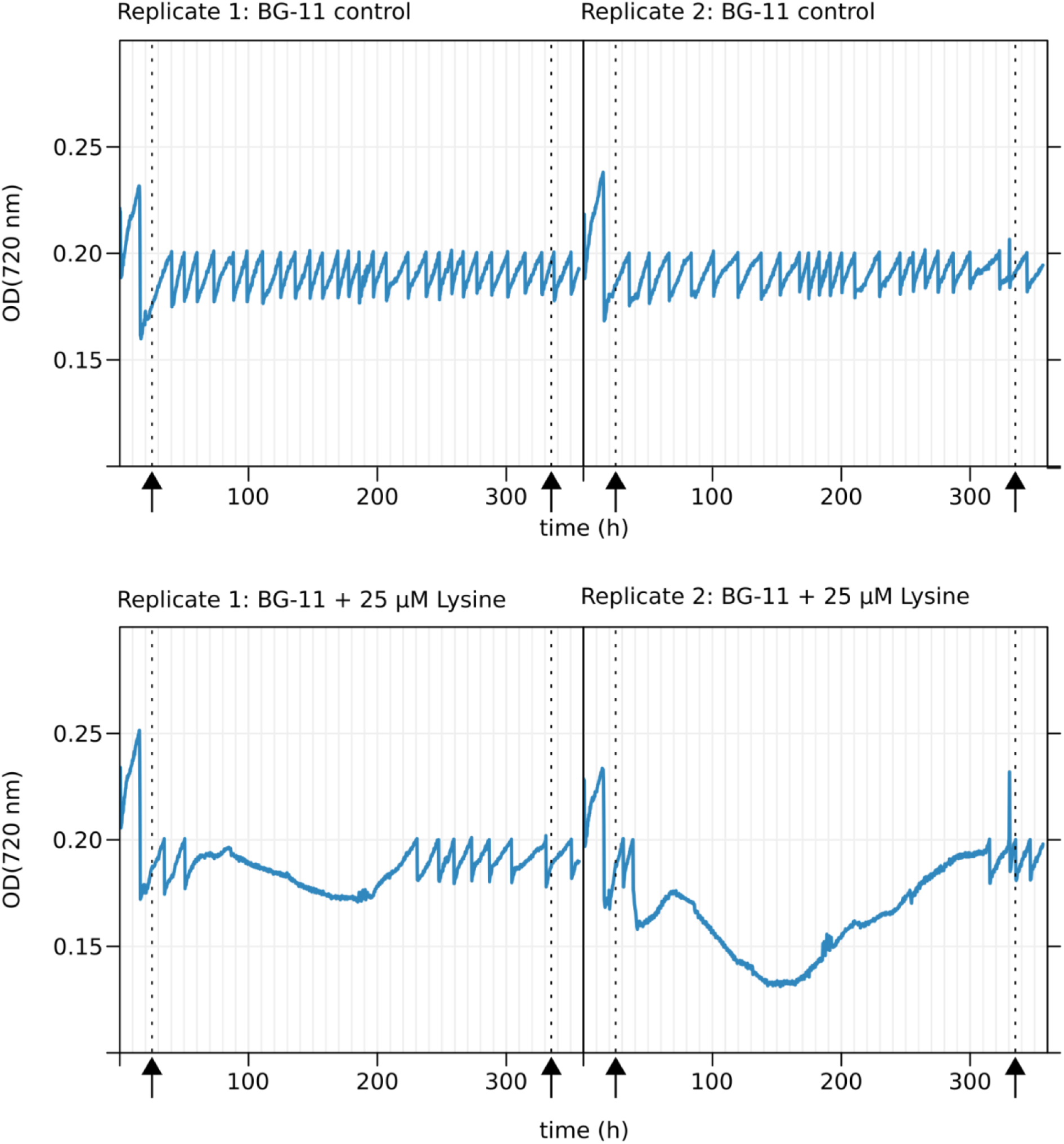
Growth of pooled CRISPRi library grown in turbidostat mode in bioreactors measured by the optical density at 720 nm. Arrows and dashed lines indicate approximate time points when samples for sequencing were taken.

**Suppl. Fig. S7**

**Supplementary Figure S7:**
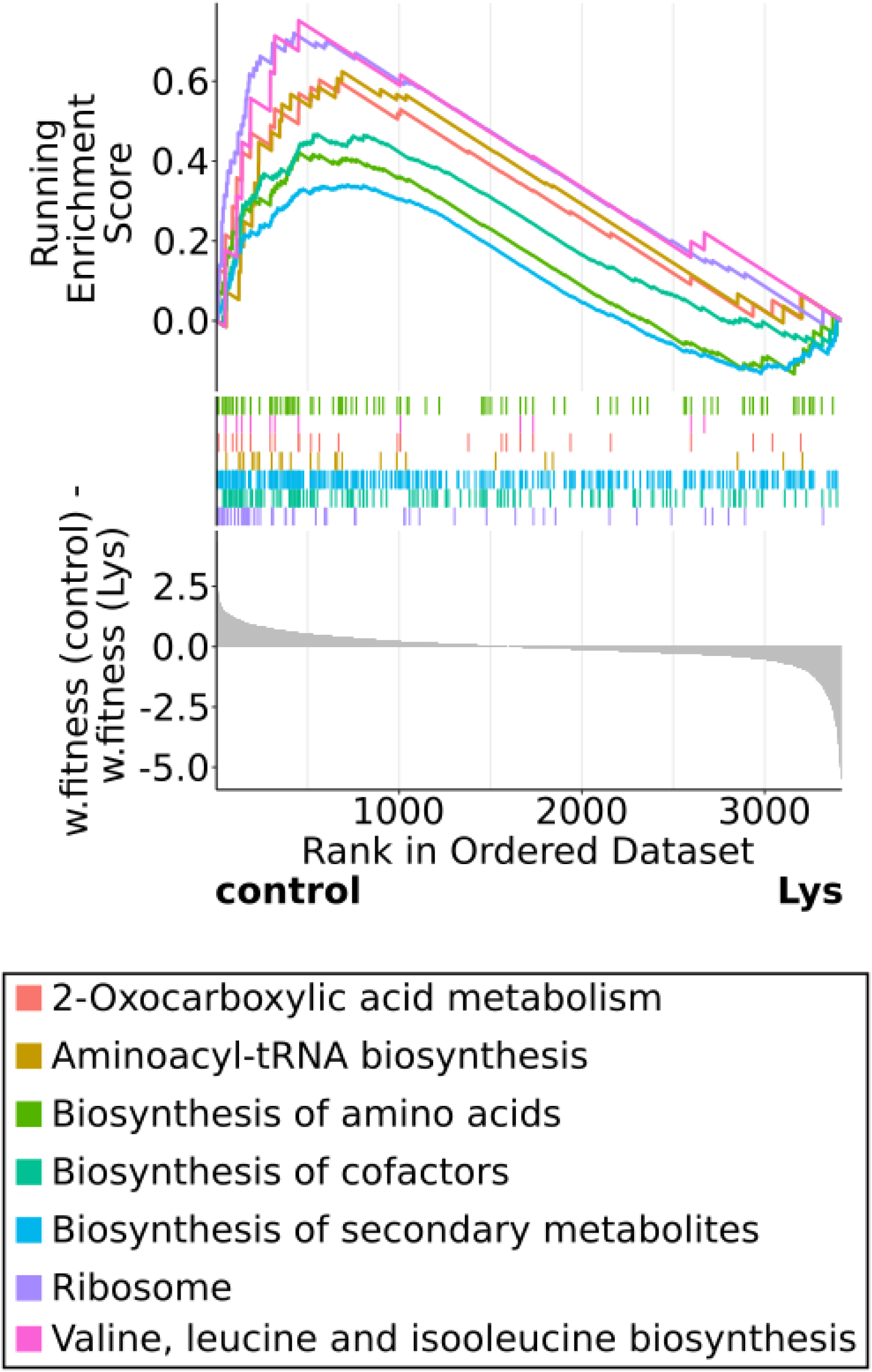
Gene set enrichment analysis (GSEA) comparing enrichment of KEGG pathways in the control library or the Lys-treated CRISPRi library. Weighted fitness values for individual genes from the Lys-treated library were subtracted from the respective values for the control CRISPRi library. The resulting score was used to rank genes, as depicted in the lowest part of the diagram. KEGG pathways are color-coded. The middle part of the diagram illustrates where genes belonging to a specific KEGG pathway are located within the ranked list. The upper most part shows the running enrichment score, i.e. the enrichment of a pathway compared to a random distribution over the whole ranked list. Compare Supp. Table S10.

**Suppl. Fig. S8**

**Supplementary Figure S8:**
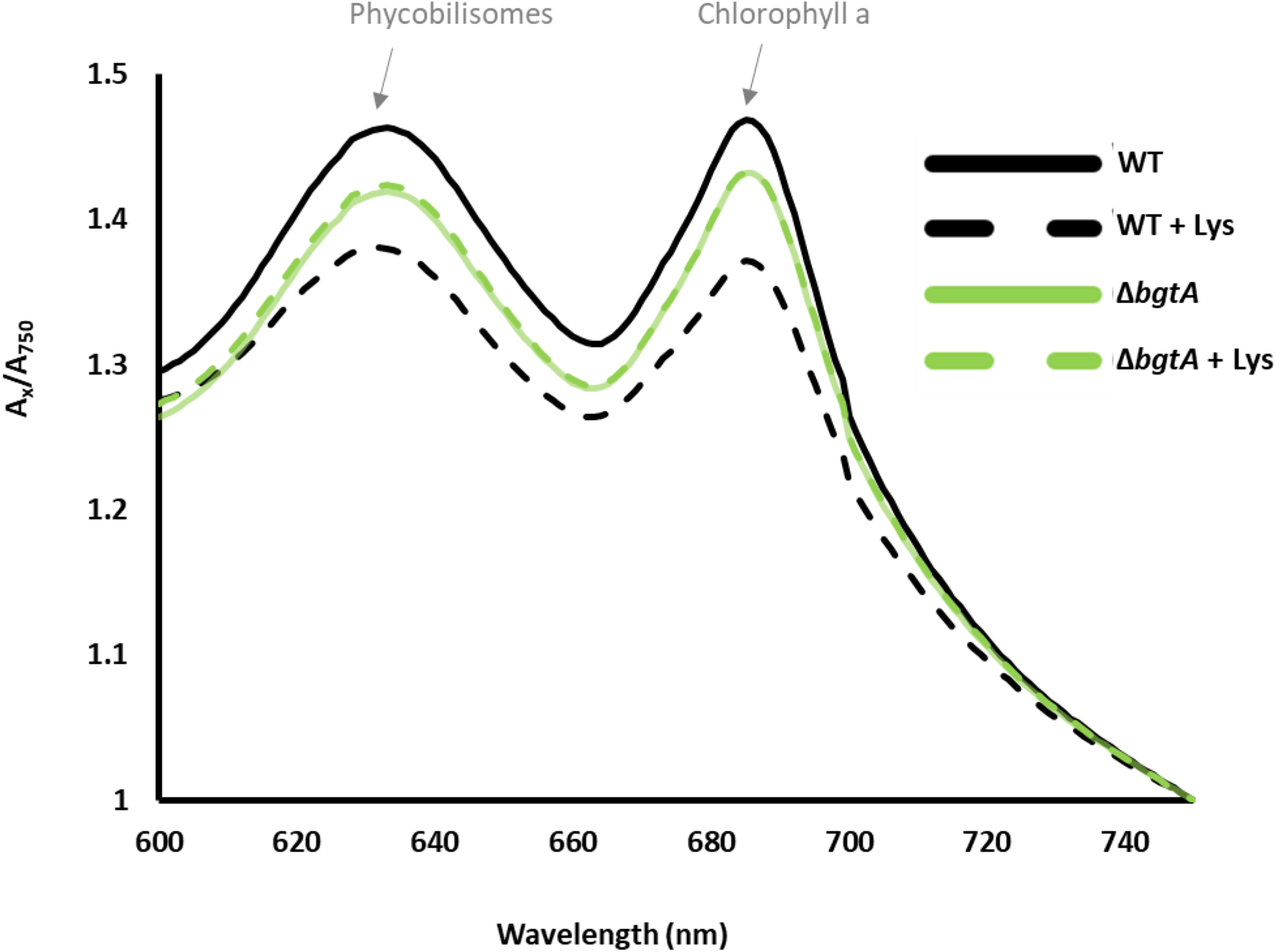
Whole-cell spectrum (WCS) of WT and Δ*bgtA* in BG11^N^ with and without 25 µM Lys. The WCS of one replicate is shown for each strain at time point 96 h. Absorbance values are normalized to absorbance at 750 nm. Absorption peaks characteristic of phycobilisomes and chlorophyll a are indicated.

**Suppl. Table S1**

**Supplementary Table S1:**
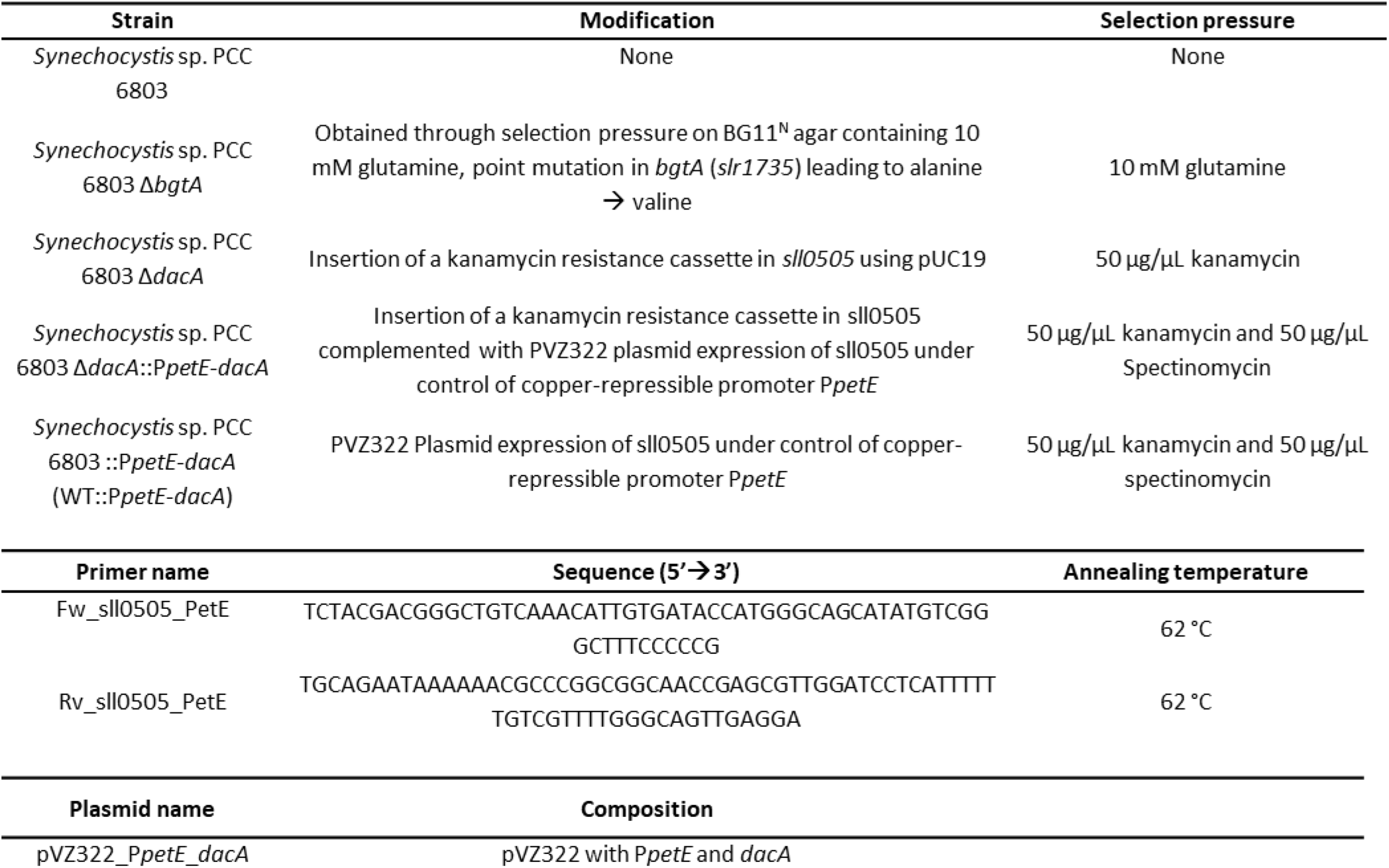
Bacterial strains, primers, and plasmids used in this study.

**Suppl. Table S2**

**Supplementary Table S2:** Chemical composition of BG11^N^ medium (modified from Rippka *et al*., 1979).

| Chemical compound | Final concentration |
| --- | --- |
| NaNO<sub>3</sub> | 17.65 mM |
| K<sub>2</sub>HPO<sub>4</sub> | 175.3 μM |
| MgSO<sub>4</sub> x 7 H<sub>2</sub>O | 304.3 μM |
| CaCl<sub>2</sub> x 2 H<sub>2</sub>O | 244.8 μM |
| Citric acid | 31.2 μM |
| Ferric (III) citrate | 24.5 μM |
| EDTA | 3.42 μM |
| Na<sub>2</sub>CO<sub>3</sub> | 377.4 μM |
| NaHCO<sub>3</sub> | 420.0 μM |
| H<sub>3</sub>BO<sub>3</sub> | 46.3 μM |
| MnCl<sub>2</sub> x 4 H<sub>2</sub>O | 9.2 μM |
| ZnSO<sub>4</sub> x 7 H<sub>2</sub>O | 0.77 μM |
| (NH<sub>4</sub>)<sub>2</sub>SO<sub>4</sub> | 1.6 μM |
| CuSO<sub>4</sub> x 5 H<sub>2</sub>O | 0.32 μM |
| Co(NO<sub>3</sub>)<sub>2</sub> x 6 H<sub>2</sub>O | 0.267 μM |

**Suppl. Table S3-10**

**Supplementary Tables S3 to S10 are available under the following link:** https://github.com/ute-hoffmann/Lys_Synechocystis/tree/main/supp_Tables

**Supp. Table S3: Overview of all sgRNAs present in CRISPRi library and associated statistical data in different conditions.** Column “time” gives the sampling point with 0 referring to the samples taken before aTc induction, time point to the second sampling point (compare Supp. Fig. S6). For a more thorough explanation of the output columns, compare https://mpusp.github.io/nf-core-crispriscreen/output. Briefly, columns “baseMean”, “log2FoldChange”, “lfcSE”, “stat”, “pvalue”, “padj” are derived from DESeq2 analyses. The remaining columns give statistics calculated to summarize the effect of different sgRNAs, as described previously (Miao & Jahn et al., 2023).

**Supp. Table S4: Data for single sgRNAs from Supp. Table S3 summarized for single genes.** “fitness_difference (control – Lys)” gives the fitness difference between the Lys-treated condition and the control condition. sgRNA targets with large absolute values in this comparison are of interest as potentially involved in conveying Lys resistance or sensitivity.

**Supp. Table S5: Gene set enrichment analysis (GSEA) of CRISPRi library data for control condition on basis of gene ontology (GO) terms.**

**Supp. Table S6: Gene set enrichment analysis (GSEA) of Lys-treated CRISPRi library on basis of gene ontology (GO) terms.**

**Supp. Table S7: Gene set enrichment analysis (GSEA) of CRISPRi library data for control condition on basis of KEGG pathways.**

**Supp. Table S8: Gene set enrichment analysis (GSEA) of Lys-treated CRISPRi library on basis of KEGG pathways.**

**Supp. Table S9: Gene set enrichment analysis (GSEA) comparing CRISPRi data for the control condition and Lys-treated cells on basis of gene ontology (GO) terms.** For this analysis, the weighted fitness values of the Lys-treated CRISPRi library was subtracted from the respective values of the control library.

**Supp. Table S10: Gene set enrichment analysis (GSEA) comparing CRISPRi data for the control condition and Lys-treated cells on basis of KEGG pathways.** For this analysis, the weighted fitness values of the Lys-treated CRISPRi library was subtracted from the respective values of the control library.

**Suppl. Table S11**

**Supplementary Table S11: Averages and standard deviations of LC-MS/MS measurements for metabolites from *Synechocystis* sp. PCC 6803 grown in BG11^N^ with and without 25 µM Ly for time points 0 h, 24 h, 48 h, 72 h, and 120 h.**

